# *Ex ante* analyses can predict natural enemy efficacy in biological control

**DOI:** 10.1101/2025.06.06.658262

**Authors:** Andrew Paul Gutierrez, Luigi Ponti, Peter Neuenschwander, John S. Yaninek, Johann U. Baumgärtner, Hans R. Herren

## Abstract

The success rate of biological control programs against invasive species is low because the efficacy of released natural enemies is unknown. Deconstruction of the successful biological control of the highly destructive cassava mealybug (CM) and cassava green mite (CGM) in Africa is used to show how *ex ante* pre-release analyses of natural enemies can increase the success rate.

A meta-population tri-trophic physiologically based demographic model (PBDM) of the cassava system accurately predicted the weather-driven daily dynamics of cassava, CM, CGM, and their introduced natural enemies across the ecological zones of Africa. Marginal analyses of the simulation data enabled parsing the differing contributions of two exotic parasitoids and a fungal pathogen in the control of CM, and the contributions of two mite predators, local pathogens and rainfall mortality in the control of CGM. The contribution of each natural enemy to the ∼95% recovery of cassava yields is estimated.

The results show *ex ante* analyses of prospective natural enemies using well parameterized PBDMs can predict natural enemy prerelease efficacy across vast geographic areas and increase the success rate of biological control programs. *Ex ante* analyses can also explain whether biological control is a viable option (e.g., Cure et al. 2020).

---

“What we observe is not nature itself, but nature exposed to our methods of questioning”.

Heisenberg (1958)

## Introduction

Since the late nineteenth century, more than 2,000 natural enemy species have been released against approximately 400 invasive species worldwide, occasionally resulting in complete control, with most introductions having limited impact (Van Lenteren *et al*., 2006; Cock *et al*., 2016). A critical question is how can the success rate be improved? To answer this, we must use *post hoc* analyses of successful programs to show why they succeeded. To this end we analyze the highly successful biological control of cassava (*Manihot esculenta* Crantz) pests across Africa, with other biological control examples reviewed in the discussion and in the supplemental materials in Gutierrez *et al*. (2025).

Cassava was introduced to Africa from Brazil during the slave trade in the 16^th^ century and became a subsistence staple crop across sub-Saharan Africa (Ferguson *et al*., 2019). Two Neotropical pests, cassava mealybug (*Phenacoccus manihoti* Matile-Ferrero (Hemiptera, Pseudococcidae))(CM) and cassava green mite (*Mononychellus tanajoa* (Bondar), (Trombidiformes, Tetranychidae)) (CGM) were accidentally introduced into Africa on cassava cutting used for plant breeding purposes early in the 1970s (Matile-Ferrero, 1977; Yaninek & Herren, 1988). The two pests quickly spread across the African cassava zone causing massive damage and creating food insecurity for >200 million Africans (Herren & Neuenschwander, 1991).

In Africa, CM was attacked by native generalist ladybird beetle predators in the genera *Hyperaspis, Exochomus* and *Diomus* but they had minimal impact (Gutierrez *et al*., 1987, 1993). CM was also infected by the endemic fungal pathogen (*Neozygites fumosa* (Zygomycetes, Entomophthorales)) (Le Rü, 1986), especially during the rainy season (Schulthess *et al*., 1987), and the effects are included in our analysis. Foreign exploration for effective adapted natural enemies was conducted in the centers of cassava origin in Central and South America that resulted in the introduction, among others, of two hymenopterous parasitoid wasps (*Anagyrus lopezi* DeSantis and *A. diversicornis* Howard) (Hymenoptera, Encyrtidae) (Herren *et al*., 1987). Both species were widely released, but *A. diversicornis* appears to have failed beyond the initial release sites in Benin (Alphen *et al*., 1990; Neuenschwander, 2001), while *A. lopezi* established widely and controlled CM populations at low levels across Africa (Neuenschwander *et al*., 1990, 1991) and in Asia where CM subsequently spread (Wyckhuys *et al*., 2018).

CGM in Africa is attacked at high densities by native generalist insect predators and endemic pathogens, but with limited impact (Yaninek & Herren, 1988). Eleven putative species of predators of CGM in the family Phytoseiidae were introduced from South America during the period 1984 to 2001, and more than 11.3 million were released in 20 African countries (Yaninek, 2007). However, only three of the predators established (*Neoseiulus idaeus* Denmark & Muma, *Amblydromalus manihoti* Moraes, and *Typhlodromalus aripo* De Leon), and only *A. manihoti* and *T. aripo* spread widely reducing CGM populations by half and increasing cassava yields by a third (Yaninek, 2007). A virulent strain of the CGM-specific fungus *Neozygites tanajoae* was also introduced from Brazil to Benin, where local impact was observed (Hountondji *et al*., 2002), but its spread was not monitored.

In this paper, we decompose the biology and dynamics of the cassava system across Africa to estimate and explain the contribution of the natural enemies in the control of CM and CGM.

## METHODS

A weather-driven physiologically based demographic models (PBDMs) of the cassava subsystems linked by dry matter and energy flow between them (Fig. 1; *cf*. Gutierrez, 1992; see Jørgensen & Nielsen, 2014) is used in our study. Only a brief review of the biology of the species in the system (Fig. 1A) is given herein, but greater details are reported in the references and summaries in the Supplemental Materials. PBDMs are time-varying life tables (Gilbert *et al*., 1976; Gutierrez, 1996) that capture the stochastic weather driven development and temporal dynamics of species independent of time and place. This property enables their use in a GIS context across vast areas such as Africa. PBDMs are bioeconomic models of species acquisition and allocation of resources (Regev *et al*., 1998; Gutierrez & Regev, 2005)

**Figure 1.**
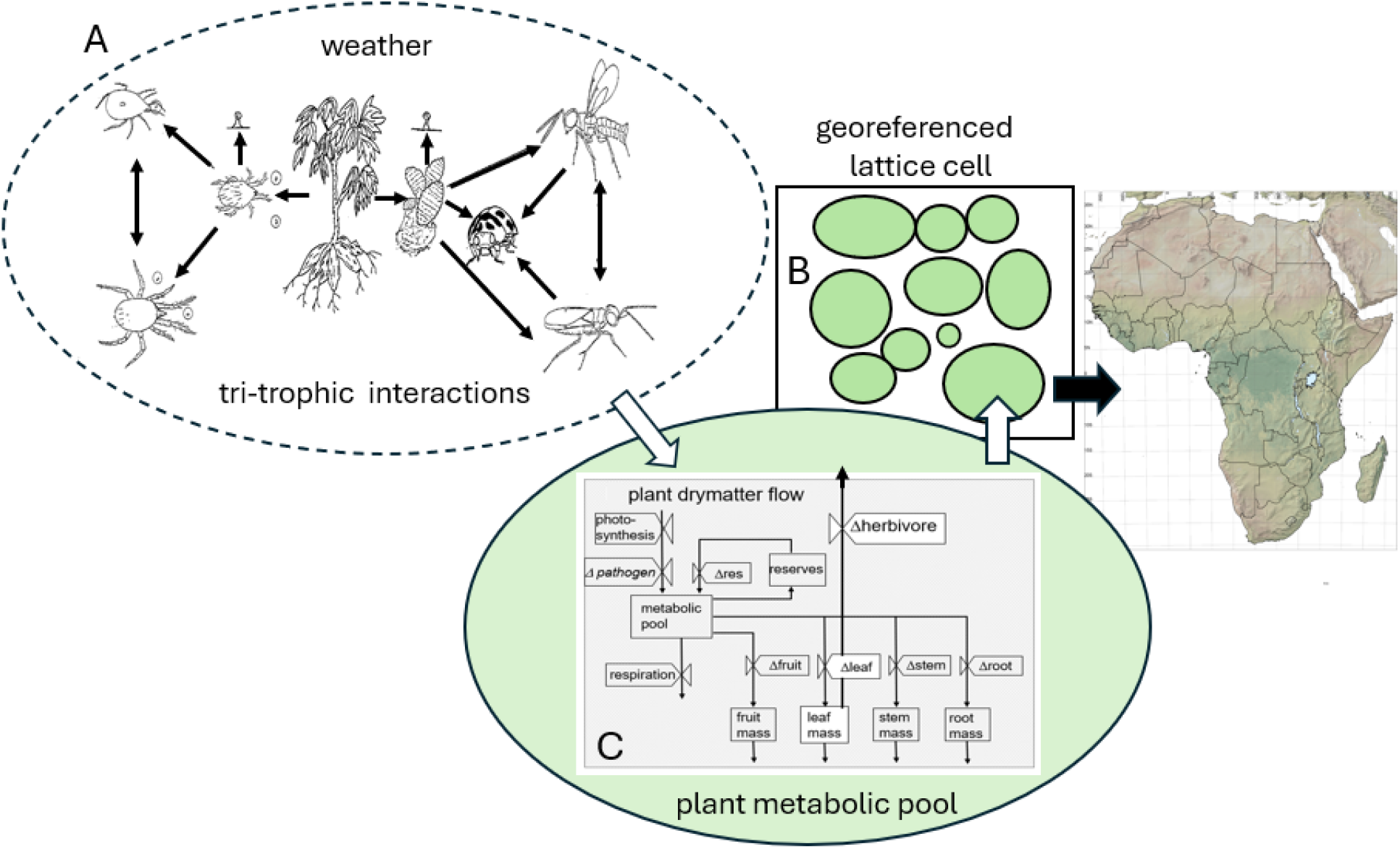
A physiologically based demographic tri-trophic meta-population cassava system for Africa: (A) tri-trophic species interactions on ten cassava plants (B) representing the dynamics the interactions depicted in Fig. 1A in each lattice cell across Africa (see text), with (C) being the within plant flow of dry matter to subunits, herbivores (i.e., CM and CGM) and higher tropic levels where single arrows indicate the direction of dry matter flow, double arrows indicating intraspecific competition, and pathogens are indicated as small symbols for conidia above CM and CGM. Individual plants in Fig. 1B have populations of the interacting species in depicted in Fig. 1A with movement of mobile arthropod stages between plants governed by species-specific resource supply/demand ratios on each plant (see Gutierrez *et al*., 1999).

The tri-trophic PBDM canopy model of the cassava system (Gutierrez, Wermelinger, *et al*., 1988; Gutierrez, Neuenschwander, *et al*., 1988; Gutierrez, Yaninek, *et al*., 1988; Gutierrez *et al*., 1993) was extended to a meta-population system composed of individual plants, each having the full complement of interacting arthropod populations (Fig. 1A; Gutierrez *et al*., 1999). Cassava variety IITA TMS30572 is used as a generic template with movement arthropods between plants modeled as a function of species-specific resource supply-demand ratio on each plant. The model algorithm accommodates a maximum of 100 plants, but to assess the daily system dynamics in each of the 40,691 ∼25 km^2^ georeferenced lattice cells across Africa on a laptop computer, ten randomly spaced plants are used. The system is modular and any combination of species with links to cassava and to each other can be introduced using Boolean variables. Biological control efforts against CM occurred during 1980-1990 and 1990-2000 for CGM, and hence the two subsystems are run separately for the two periods. The only change to the model was a nonlinear developmental rate model for cassava (Gutierrez *et al*. 1999; see Supplemental Materials). The two major components of PBDMs are: (i) the age-structured dynamics model and (ii) parameterization of the biological functions capturing the biology of the species.

### Age-structured dynamics model

Cohorts of ectotherm organisms have temperature-dependent distributed maturation times with means and variance. Further, each species/stage has different responses to temperature, resulting in different time scales for each. The time-invariant distributed maturation time dynamics model with attrition (Manetsch, 1976; see Vansickle, 1977 for the time vary form) accommodates this biology and is used to develop the age-mass structured PBDMs of the species (see appendix and Supplemental Materials). Other dynamics model may also be used (Gutierrez, 1996; Di Cola *et al*., 1999; Buffoni & Pasquali, 2007). Two approaches were used to parameterize the dynamics models: the metabolic pool (MP) and biodemographic function (BDF) approaches (see Gutierrez et al. 2025).

PBDM/MP models capture per capita biomass/energy acquisition and allocation to growth and reproduction and the effects on mortality in the plant and arthropod populations. Cassava plants search for light, water, and nutrients to produce photosynthate that is allocated to plant subunit growth and to reproduction (*cf*. de Wit & Goudriaan, 1978). Gutierrez et al. (1975) and Wang *et al*. (1977) used the MP approach to develop a PBDM canopy model of linked age-mass structured populations of leaves, stem, root and fruit. Gutierrez and Baumgärtner (1984) used MP notions to develop PBDM models for growth, development, and reproduction of arthropods. Most of the species in our study are MP based. Further, migration rates of arthropods between plant are functions resource acquisition success.

In the PBDM/BDF approach, developmental rates and birth and death rates of the species are estimated from age-specific life table studies conducted under an array of temperatures and other conditions (Gutierrez & Ponti, 2013), but field studies can also be used (e.g., Dreyer & Baumgärtner, 1995). The vital rates are the outcomes over the life cycle of a cohort of organisms of how they acquire and allocate energy, survive, and reproduce under the experimental conditions – i.e., the results of metabolic pool processes. BDFs summarize the effects of abiotic variables on species developmental, birth, and death rates, and are used to parameterize the age-structured, weather-driven models. The most common BDFs are depicted in a stylized manner in Fig. 2, but other BDFs may be developed to accommodate additional aspects of the biology of species (see Supplemental Materials).

**Figure 2.**
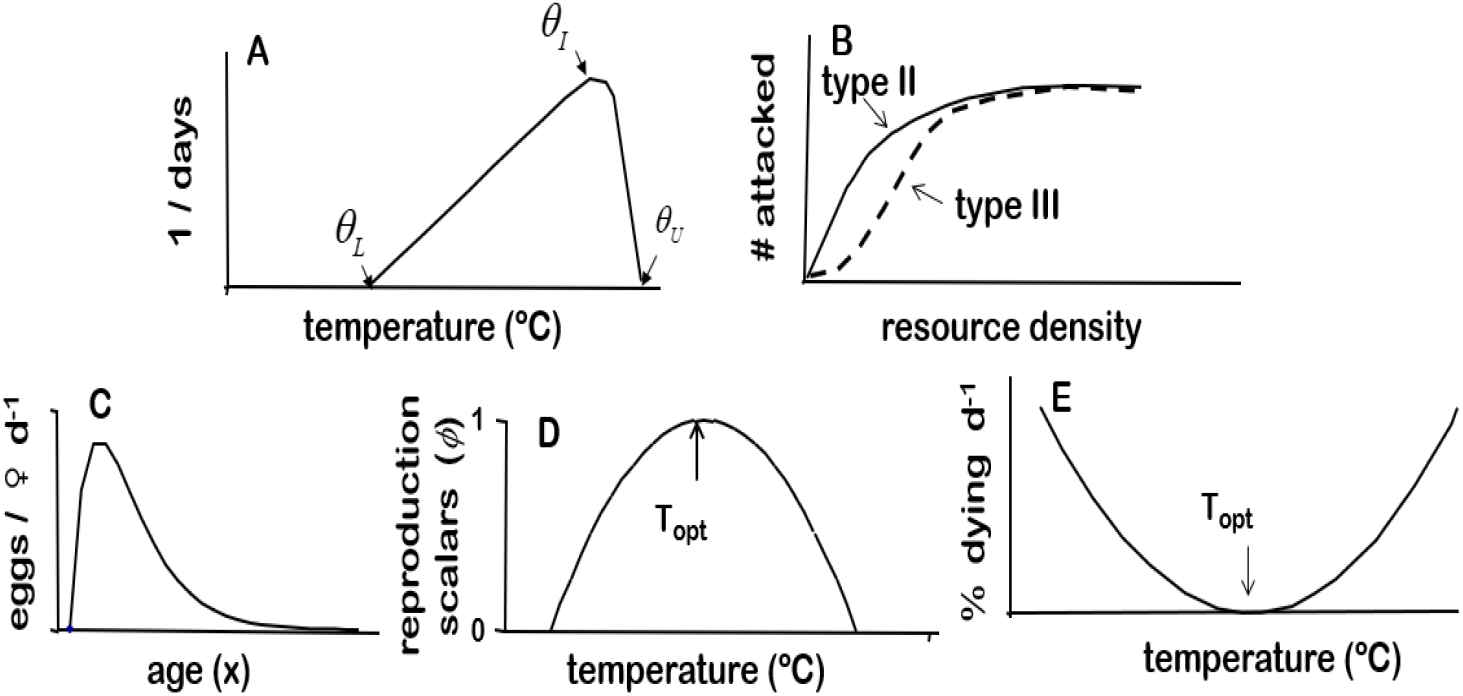
Minimum stylized biodemographic functions required to develop PBDMs (modified from Gutierrez & Ponti, 2013; Gutierrez et al. 2025): (A) rate of development on temperature, (B) functional response for resource acquisition, (C) age-specific reproductive profile at the optimum temperature, (D) temperature scalar to correct reproduction from the optimum temperature, and (E) mortality rate per day on temperature. Note that similar shaped scalars to D and E can be developed for the effects of relative humidity and other factors. In practice, the functions D and E are not symmetrical.

### Weather data

The system model is run using daily maximum and minimum temperature, precipitation, solar radiation, and relative humidity as inputs to the functions in Fig. 2 (see Supplemental Materials). Daily weather data for 40,691 ∼25 km^2^ georeferenced lattice cells across Africa for the 1980-2010 period were sourced from the AgMERRA global weather dataset created by the Agricultural Model Intercomparison and Improvement Project (AgMIP, https://agmip.org/) (Ruane *et al*., 2015). The AgMERRA data were accessed through the Goddard Institute for Space Studies (GISS) of the National Aeronautics and Space Administration (NASA, https://data.giss.nasa.gov/impacts/agmipcf/). From the perspective of ectotherms, climate change is simply another weather pattern, and future weather projections from climate models could easily be used to run the model. The effects of climate change on the system are beyond the scope of this study (see Supplemental Materials in Gutierrez et al. 2025).

For feasible daily computations over ten-year periods on a laptop computer, alternate longitude-latitude lattice cells (∼10,340) were used that provided sufficient grain to characterize the distribution and relative abundance of the species across Africa. The same nominal initial conditions were used for all lattice cells, and georeferenced annual summary simulation variables for all populations were written to text files. Excluding the first year data when the system is assumed equilibrating to lattice cell weather, means, standard deviations, and coefficients of variation of all annual summary variables are computed across years for each cell and used for GIS mapping and for statistical and marginal analyses.

### GIS mapping

The open source GIS software GRASS (Neteler *et al*., 2012; GRASS Development Team, 2022) (http://grass.osgeo.org/) was used to map the summary output data. The current GIS distribution of cassava cultivation in Africa use in our analyses is from Tang *et al*. (2024; CROPGRIDS). Other GIS data layers used in mapping were sourced from the public domain repository *Natural Earth* (http://www.naturalearthdata.com/)). Inverse distance weighting bicubic spline interpolation was used to create a continuous raster surface of the simulation data.

### Marginal analysis

A general binomial linear regression model (eqn. 1) is used to summarize the Africa-wide simulation data using species absence-presence as independent dummy variables (i.e.,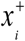, [0, 1]) with dependent variables being root dry matter yield or pest density (*Y*),

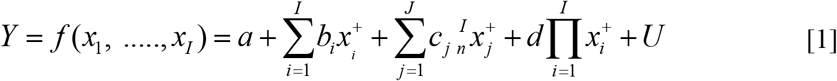

where 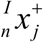 is the *j*^*th*^ interaction of the 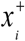 taken *n* at a time, 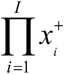 is the product of the *I* variables, and *U* is the error term. Only significant terms (*p* < 0.05) are included in the final model. The marginal impact of species presence 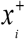 on say cassava yield (g dry weight) or pest density is estimated as 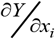 given the average effects of the other independent variables.

## RESULTS

### Cassava

The distribution of simulated prospective average root yields is mapped in Fig. 3A for the period 1981-1990, while the same data masked for the known distribution of cassava (Tang *et al*., 2024) are mapped in Fig. 3B. Average annual degree days computed using a nonlinear model (*dd>*14.86°C) for cassava (Supplemental Materials eqn. 1) are mapped in Fig. 3C, and average annual rainfall mapped in Fig. 3D with the limits of cassava production indicated by dashed white lines. Note that the simulated distribution of cassava coincides well with its known distribution above the ∼700 mm rainfall isocline. The model predicts yield in South Sudan and parts of Uganda, Kenya, and Ethiopia (Fig. 3A vs masked Fig. 3B) where alternative crops (e.g., maize, sorghum) are more important. Difference in distribution are also due to the use of observed weather in the simulation (e.g., Fig. 3D), while Tang *et al*. (2024) report an aggregate of historical records. All subsequent results are masked for the distribution of cassava.

**Figure 3.**
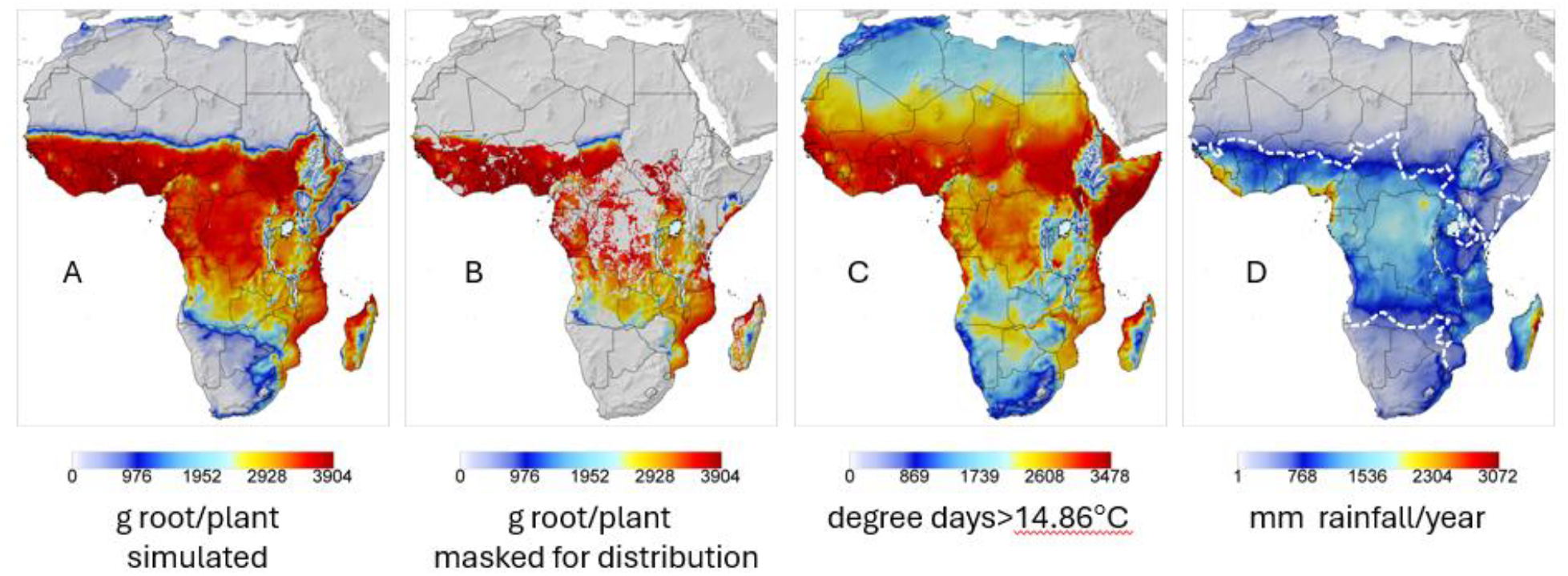
Cassava per plant averages for years 1981-1990: (A) average simulated prospective distribution and relative root yield (g dry matter) in Africa, (B) root yield from masked for the known distribution of cassava in sub-Saharan Africa (Tang *et al*., 2024) (public repository: https://doi.org/10.6084/m9.figshare.22491997), (C) average annual degree days >14.85°C for cassava estimated using a nonlinear model (Supplemental Materials eqn. 1) based on cassava growth rates data (Keating & Evenson, 1979), and (D) mean annual mm rainfall with the distribution of cassava circumscribed by the ∼700 mm rainfall isohyet (dashed white lines).

### Cassava mealybug

Using 30/6/1982 to 30/6/1985 weather at Cotonou, Benin, West Africa and given the action of *A. lopezi, A. diversicornis* and the endemic fungal pathogen, rapid control of CM is predicted (Fig. 4A). Data for Ibadan, Nigeria (Gutierrez, Neuenschwander, *et al*., 1988) are illustrated as an inset in Fig. 4A. Prospectively, *A. lopezi* in concert with rainfall/fungal mortality rapidly suppresses CM populations to very low levels causing near total displacement of *A. diversicornis* (Gutierrez *et al*., 1999). However, absent *A. lopezi*, high populations of *A. diversicornis* are predicted that do not control CM (Fig. 4B). Although *A. diversicornis* appears to play a minor role in suppressing CM (see Neuenschwander, 2001), it is included in the Africa-wide study to explore its potential in the other ecological regions.

**Figure 4.**
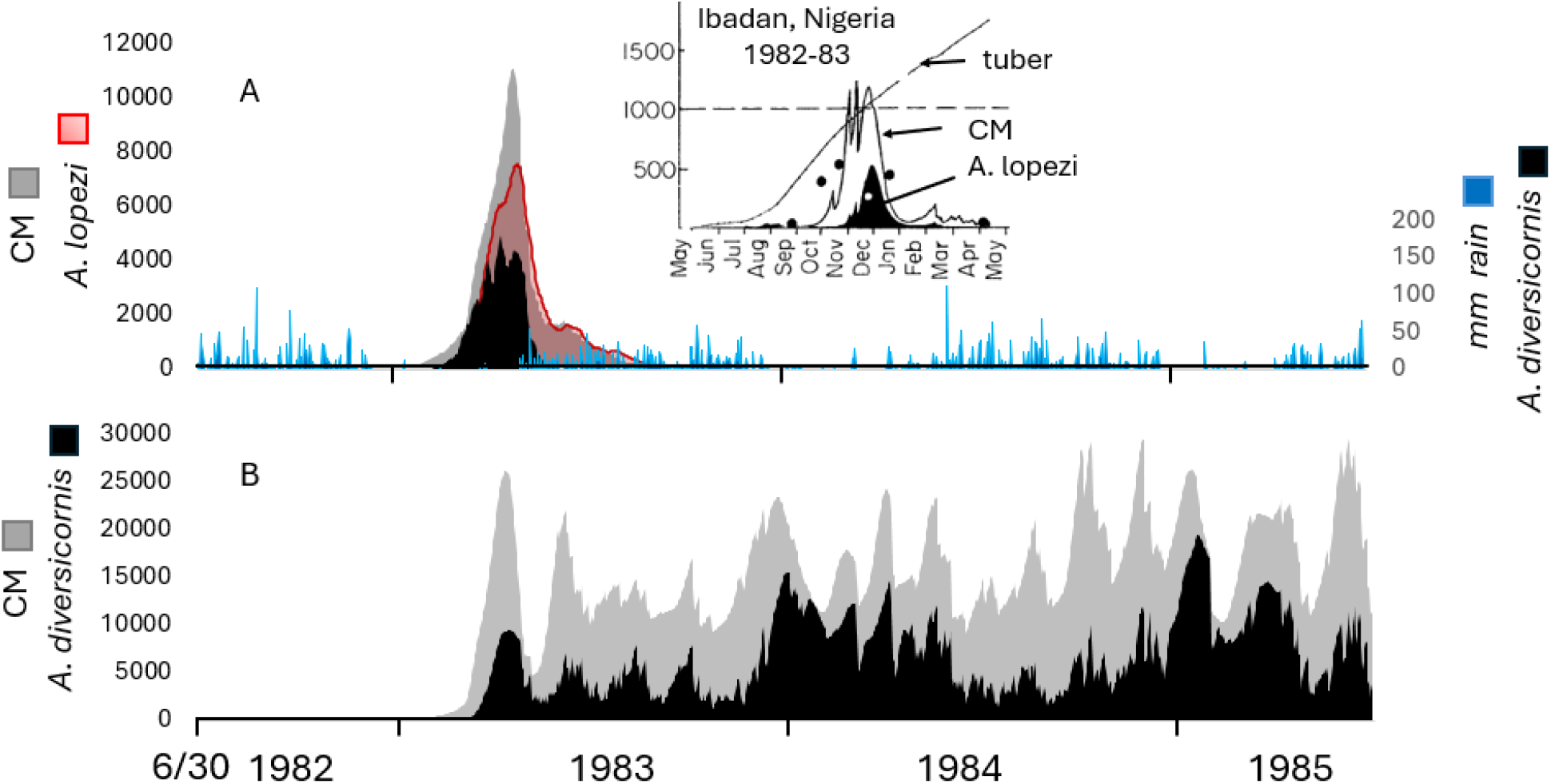
Simulated average per plant population dynamics at Cotonou, Benin during 6/30/1982 to 6/30/1985: (A) cassava mealybug, and CM parasitized by *A. lopezi* and *A. diversicornis* given the impact of fungal pathogen mortality on CM, and (B) CM and *A. diversicornis* parasitized CM given fungal mortlity. Daily rainfall is indicated as the vertical blue lines in 4A, and the inset is from Gutierrez *et al*. (1988).

Absent natural controls, simulated average annual root yields (g dry weight/plant) over the 1981-1990 period are mapped in Fig. 5A with CM densities mapped as log10 average CM active stages in Fig. 5B. Prospective root yield losses are mapped in Fig. 5C as the difference of data in Fig. 3B and Fig. 5A. Predictions of the geographic distribution of CM (Fig. 5B) agrees qualitatively with the predictions of Yonow *et al*. (2017; inset) using the species distribution model (SDM) CLIMEX.

**Figure 5.**
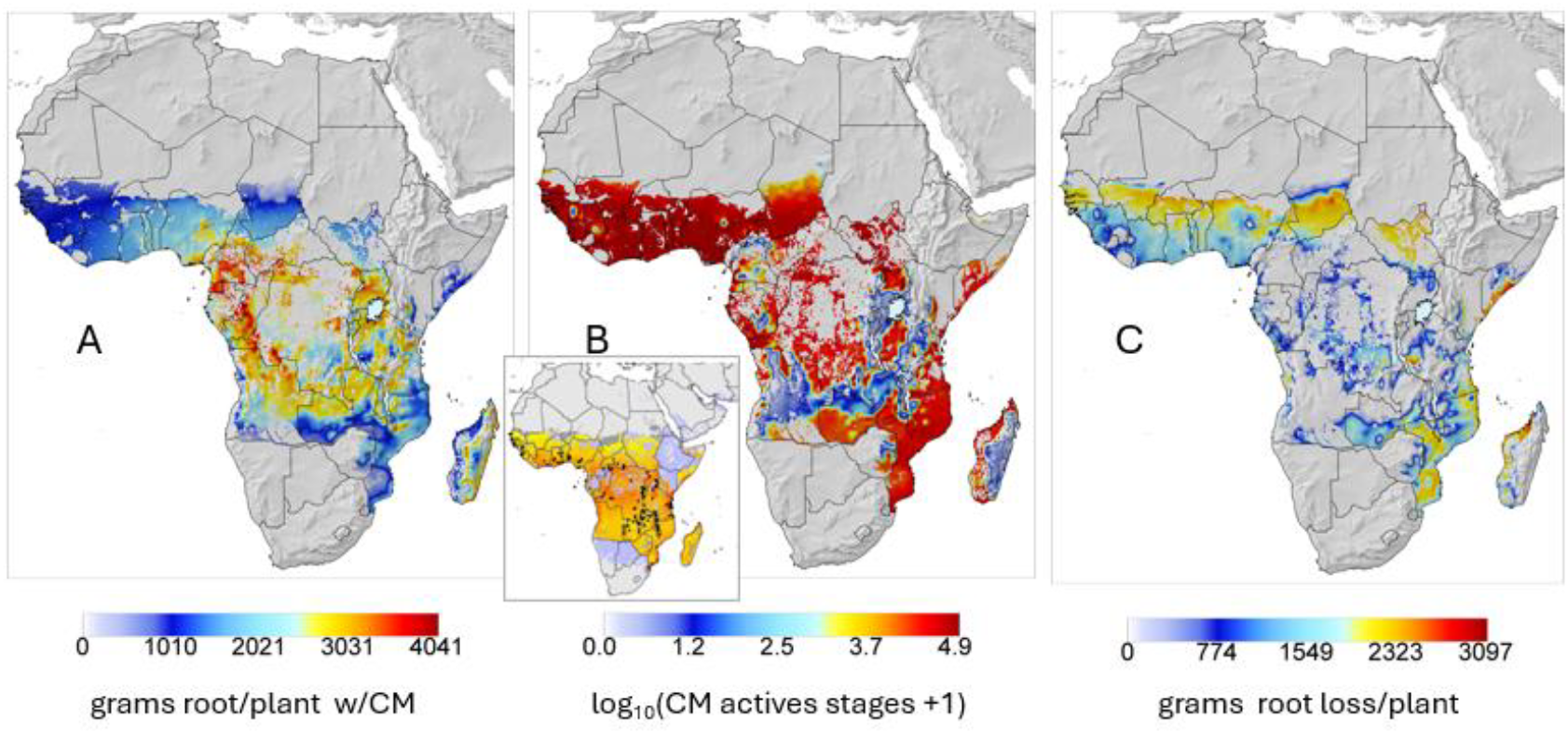
Prospective averages over the 1981-1990 period before successful biological control of CM given the effect of the fungal pathogen on CM: (A) prospective simulated distribution of cassava root yield (g dry matter) after the invasion of the cassava mealybug (CM), (B) prospective distribution of log10 mealybug active stages with an inset clip of the predicted CLIMEX distribution of CM by Yonow *et al*. (2017), and (C) yield loss due to uncontrolled mealybug computed as yields in Fig. 3A minus yields in Fig. 5A. The small difference in the scales between Figs. 3A and 5A are due to GIS system interpolation estimates.

With the control of CM (Fig. 5B vs Fig. 6B) principally by *A. lopezi*, prospective cassava yields rebounded to ∼95% of pre-mealybug invasion levels (Fig. 5A vs Fig. 6A). The distribution and relative average cumulative densities of *A. lopezi* and *A. diversicornis* after control of CM are mapped in Figs. 6C,D respectively showing the distribution of *A. lopezi* is wider than that of *A. diversicornis*. Highest CM post control densities are predicted in the drier northern and eastern areas where cassava production is marginal, and the effects of rainfall on pathogen activity are reduced (see discussion).

**Figure 6.**
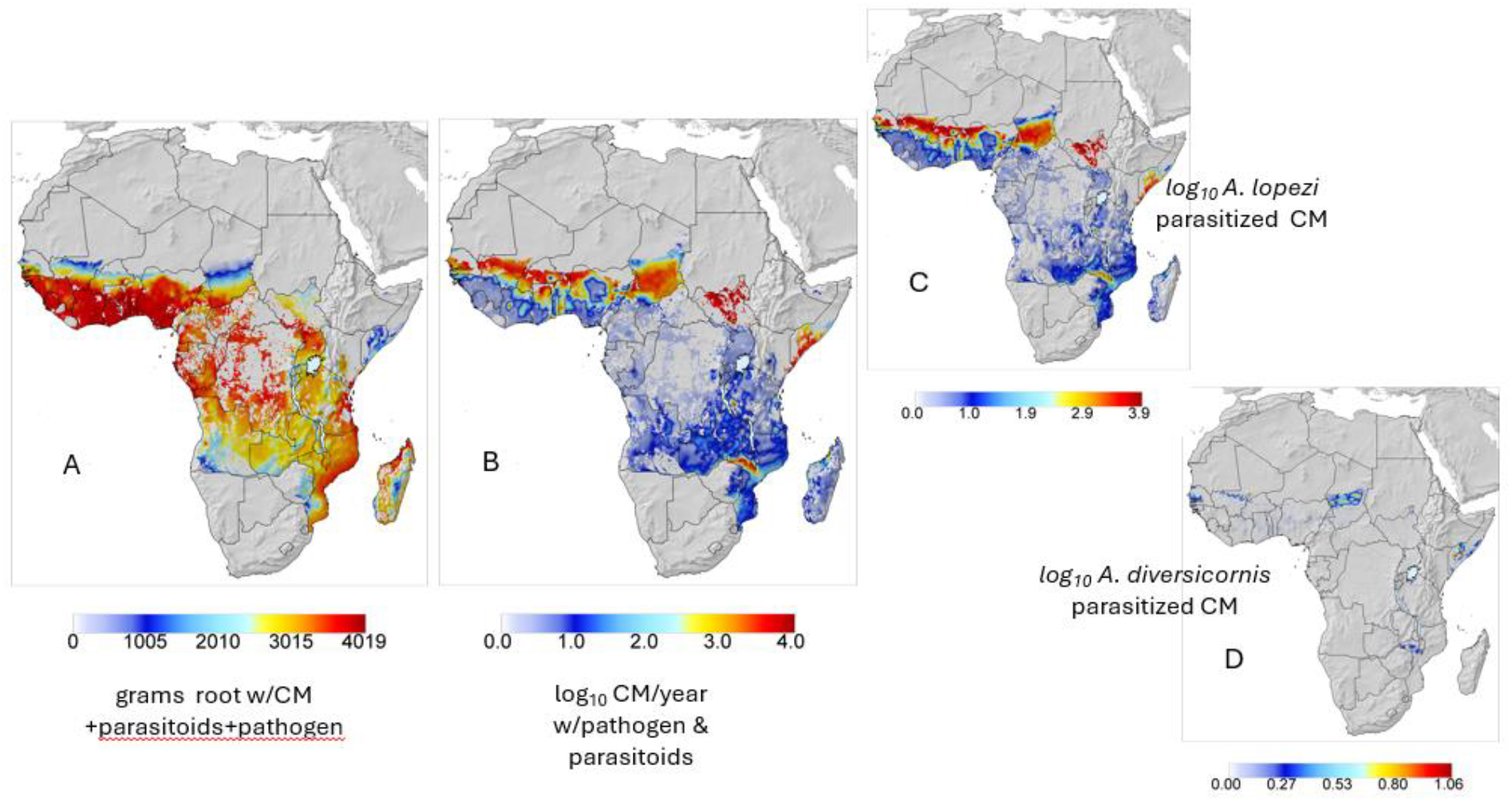
Prospective annual averages (1981-1990) after the release of *A. lopezi* and *A. diversicornis*: (A) relative root yield (g dry matter), (B) annual cumulative log10 CM active stages, (C) log10 *A. lopezi* parasitized mealybugs, and (D) log10 *A. diversicornis* parasitized mealybugs. The effect of the endemic fungal pathogen on CM is included in all sub figures. The results are masked for the known distribution of cassava (Tang *et al*., 2024).

### Marginal analysis of cassava yield and control of CM

Data from lattice cells with root yields greater than 1,500g (i.e., hereafter called the cassava belt) were used in this and subsequent analyses (eqn. 2). *Y* is average gram root dry matter/plant and the arthropod species presence variables 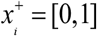) are mealybug (*CM*^*+*^), *A. lopezi* (*Al*^*+*^), *A. diversicornis* (*Ad*^*+*^), and pathogen (*P*^*+*^).

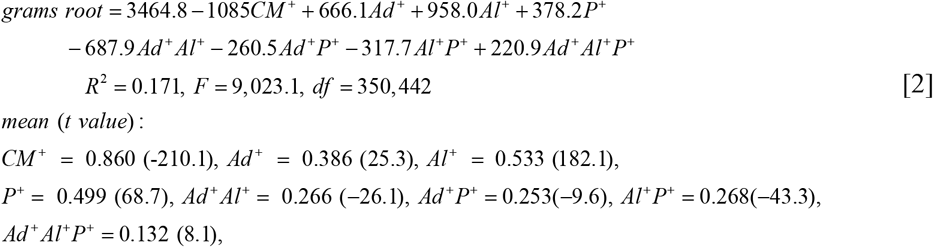

The marginal effect on average grams root dry matter loss/plant due to CM is 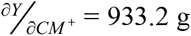, while the average marginal contributions to yield increase are 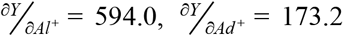, and 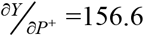 respectively for a combined total compensation of 923.8 g. The apparent positive contribution of *A. diversicornis* to yield occurs because *Al*^*+*^ was absent in numerous runs, and *Ad*^*+*^ partially suppresses *CM* (e.g., Fig. 4B). However, given *Al*^*+*^, the expected contribution of *Ad*^*+*^ is low (Fig. 6D). Excluding *Ad*^*+*^ from the analysis gives eqn. 3.

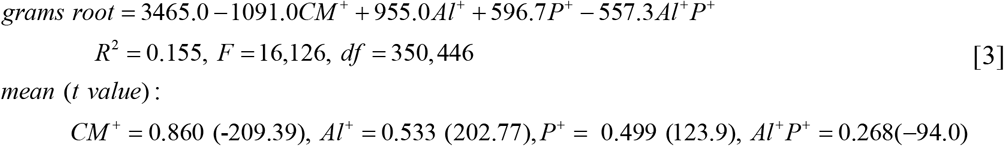

Substituting average values, 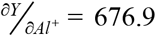 and 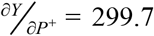 for a combined marginal compensation to yield of 976.6 g, which is roughly the same as from eqn. 2. The contribution of *Al* ^+^ to yield is ∼2.25-fold that of *P*^+^ supporting field observations (Hennessey *et al*., 1990; Neuenschwander *et al*., 1989, 1990, 1991). The suppressive effect of the pathogen on CM is likely an overestimate, but this does not detract from the overall conclusions.

### Control of CM

Using cumulative daily CM plant^-1^ year^-1^ (i.e., CM days) as the dependent variable, with *Al* ^+^ and *P*^+^ as independent variables give eqn. 4.

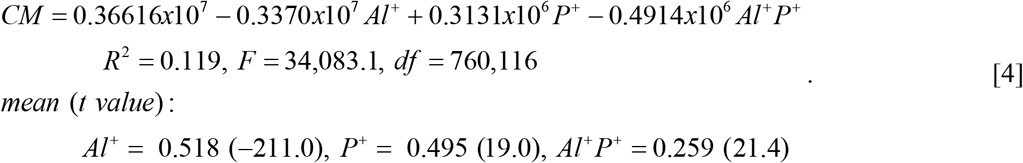

Using mean values, 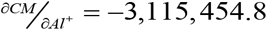 and 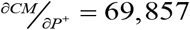 shows that the negative effect of *Al*^*+*^ is large and reduces CM density, while the positive contribution of *P*^*+*^ is the net effect of precipitation on pathogen mortality to CM and its enhancement of plant growth that increases CM. The combined action of *Al*^*+*^ and *P*^*+*^ reduces average cumulative CM active stages ∼87%.

### Cassava green mite

The simulated dynamics of the CGM/predator system at the initial release site of Ikpinlè, Benin during 6/1/1990 to 6/30/2000 are summarized in Figure 7. Before the introduction of the predators, CGM mortality largely accrued from the interaction of rainfall and endemic fungal pathogens (Fig. 7A,B; Yaninek *et al*., 1989, 1996). With the introduction of mite predators, highest CGM densities occurred during the dry season (Fig. 7B; see Yaninek *et al*., 1987) with lower densities due to the action *T. aripo* and *A. manihoti* (Fig. 7C, D). The simulation results support observations that *T. aripo* is the most effective predator (Yaninek *et al*., 1998; Yaninek, 2007). Monthly estimates of *CGM* and *T. aripo* densities on thirty plants during 1994-1997 at Ikpinlè, Benin confirm persistence of the *CGM* /*T. aripo* interaction over multiple seasons (Fig. 7E).

**Figure 7.**
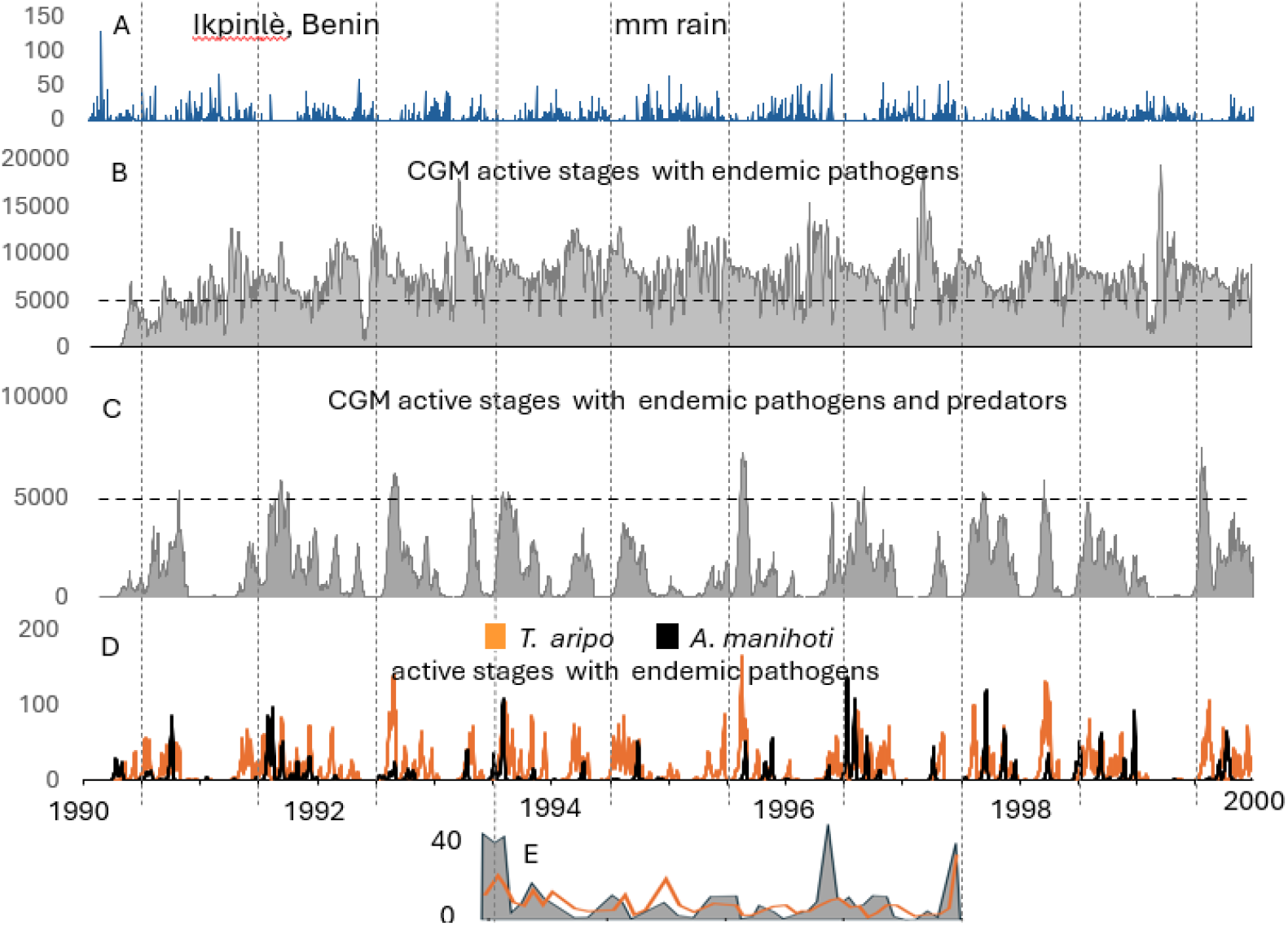
Simulated population dynamics of CGM and introduced predators with the combined action of rainfall and endemic pathogens using weather data from 6/1/1990 to 6/30/2000 for the lattice cell with Ikpinlè, Benin: (A) mm of daily rainfall, (B) CGM active stages absent predation, and (C) CGM with *T. aripo* and *A. manihoti*, (D) the dynamics of *T. aripo* and *A. manihoti* active stages, and (E) on-farm observations on 30 plants at approximately monthly intervals of average active stage counts of CGM (grey area) on the first developed leaf and *T. aripo* (orange line) in the shoot tips (*T. aripo* = 0.457*CGM*, R ^2^ = 0.675 ; Yaninek, unpub. data). The horizontal lines at 5000 in B and C are reference lines.

### Marginal analysis of yield with and without CGM

Average annual rainfall during 1991-2000 is mapped in Fig. 8A, and prospective average cassava yields absent CGM are mapped in Fig. 8B. Given CGM and the action of endemic fungal pathogens, prospective cassava yields losses range from ∼15% to ∼67% (Fig. 8B vs Fig. 8C) which agree with measured yield losses of 13 to 80% (Yaninek *et al*., 1990). Further, the prospective distribution of CGM (Fig. 8E) is comparable to that predicted by Parsa *et al*. (2015) using the SDM MaxEnt algorithm (e.g., Phillips & Dudik, 2008).

**Figure 8.**
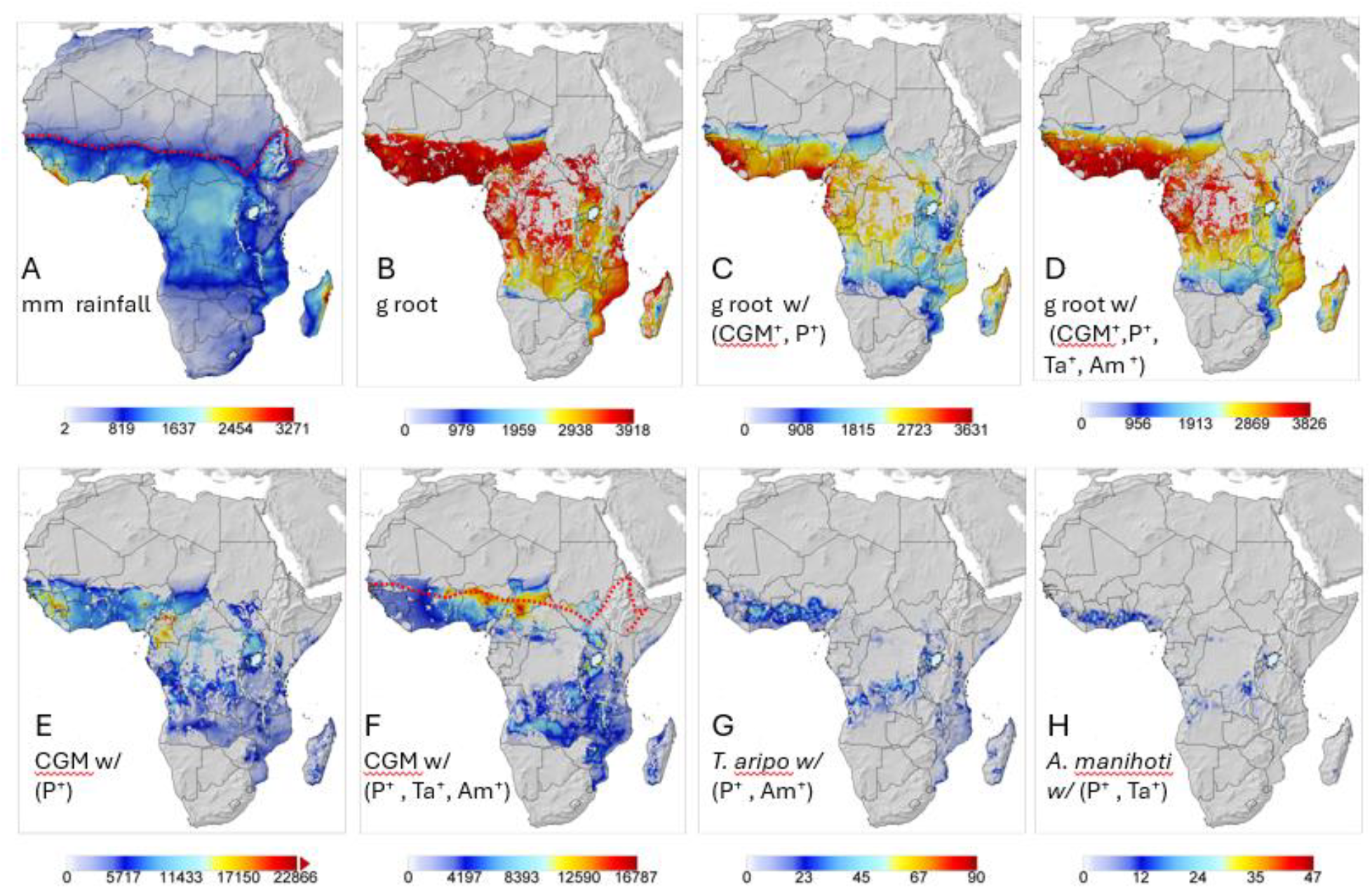
Annual averages using 1991-2000 weather data: (A) average rainfall, (B) prospective relative root yield (g dry matter) absent CGM, (C) root g dry matter with CGM^+^ and endemic diseases *P*^*+*^, (D) g root with CGM^+^, endemic diseases P^+^, *T. aripo* (*Ta*^*+*^), and *A. manihoti* (*Am*^*+*^), cumulative values of (E) CGM active stages with *P*^*+*^, (F) CGM active stages with *P*^*+*^, *Ta*^*+*^ and *Am*^*+*^, (G) *T. aripo* active stages given *P*^*+*^ and *Am*^*+*^, and (H) *A. manihoti* active stages given *P*^*+*^ and *Ta*^*+*^. The simulation data are masked for the distribution of cassava production (Tang *et al*., 2024) and the red dashed line subfigure F is the northern ∼800 mm rainfall isohyet from subfigure 8A.

The introduction of the mite predators increased prospective root yields to ∼ 95% of potential (Fig. 8B vs Fig. 8D) due to control of CGM (Fig. 8E vs Fig. 8F). Average densities of the predators are summarized in Fig. 8G, H showing *T. aripo* has a wider distribution and its maximum densities are ∼1.5-2.0 fold greater than *A. manihoti*.

### Marginal analysis of cassava yield and control of CGM

**Cassava yield** - Using lattice cell data across the cassava belt and absence/presence variables for (*CGM*^*+*^), *T. aripo* (*Ta*^*+*^), *A. manihoti* (*Am*^*+*^), and pathogen (*P*^*+*^) gives regression eqn. 5.

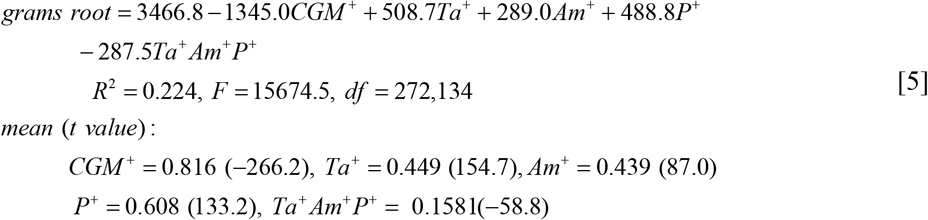

Using mean values, average root loss 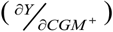 due to CGM in the absence of natural controls is - 1,097.5 g (31.6% loss). The order of factors increasing yield is *P*^*+*^> *Ta*^*+*^*>Am*^*+*^, and their combined action reduces CGM damage by more than 55.1% (i.e., 604.2 g) for an average 14.2% root loss. The effect of *P*^*+*^ is the net of pathogen mortality to CGM and the enhancing effects of rain on root yields. The interaction term *Ta*^*+*^*Am*^*+*^*P*^*+*^ is negative suggesting an adverse effect of interspecific competition but resulting in only a 45.4 g (∼1.3%) reduction in average root yield.

### Control of CGM

Using annual cumulative CGM eggs plus immature stages/plant as the dependent variable and *Ta*^*+*^, *Am*^*+*^, *P*^*+*^ as independent variables gives eqn. 6.

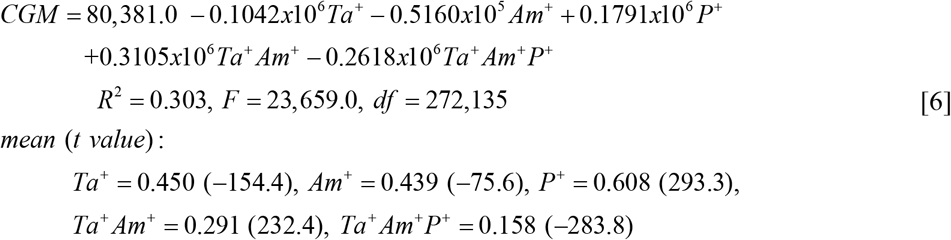

The coefficients of *Ta*^*+*^, *Am*^*+*^ and *Ta*^*+*^*Am*^*+*^*P*^*+*^ are negative indicating they reduce CGM levels, while the interaction term *Ta*^*+*^*Am*^*+*^ is positive and is a metric of the effect of interspecific competition. The coefficient of *P*^*+*^ is positive due to the net effect of rainfall/pathogen mortality on *CGM* and rainfall that favors plant and CGM growth (Fig. 7B vs 7C). Using mean values, the marginal effects are: 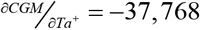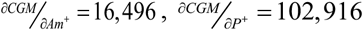. Across the three factors, the average net annual cumulative CGM active stages is 162,025. Removing *P*^*+*^ to assess the action of the two predators yields eqn. 7.

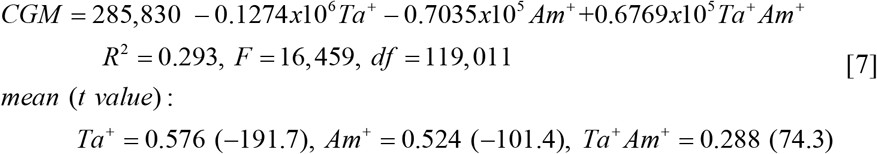

The impact of *Ta*^*+*^ 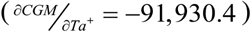 is ∼3-fold that of *Am*^*+*^ 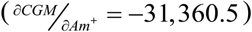, and the average net annual cumulative CGM stages is 162,539 (eqn. 7) vs 162,025 (eqn. 6) suggesting a very small irreplaceable mortality from *P*^*+*^ of ∼0.3%. On average, the predators reduce prospective CGM populations by 56.7%.

## DISCUSSION

Worldwide damage from invasive species since 1960 has been US$ 1,130.6 billion (Cuthbert *et al*. 2022) with annual losses in the USA of more than US$ 140 billion (Pimmentel *et al*. 2000), and annual losses in the European Union are US$ 28 billion with projected losses of US$148 billion by 2040 (Henry *et al*. 2023). Loss of subsistence staple food crops (e.g., cassava, maize, millet, sorghum) and in veterinary and human health in Africa and other lesser developed areas are large but not well documented (Pyšek et al. 2008; Khuroo *et al*. 2011). Classical biological control programs against invasive species can be highly effective (*cf*. Huffaker & Messenger, 1976), but they have a very high failure rate (Van Lenteren *et al*., 2006; Cock *et al*., 2016). A contributing factor in the failures is that target invasive species may have wide native ranges with physiological strains adapted to local conditions (e.g., Gutierrez *et al*., 2025b) and associated adaptation in their natural enemies making discovery of efficacious natural enemies vexing (e.g., alfalfa weevil (*Hypera postica* (Gyllenhal)), Salt & van den Bosch, 1967; walnut aphid (*Chromaphis juglandicola* (Kale)), van den Bosch *et al*., 1979).

The biological control of cassava pests in Africa is an exemplar of biological control of the highly destructive cassava mealybug (CM) and cassava green mite (CGM) (Neuenschwander *et al*., 2023) that kept food insecurity at bay for > 200 million people (Neuenschwander *et al*., 1989). Depending on monetary scenarios, the economic gains from the control of CM alone ranged from US$ 10 - 30 billion, with benefit-cost ratios of 200 (Norgaard, 1988) and 370-740 (Zeddies *et al*., 2001; Neuenschwander, 2004). The cost of the biological control program for CM and CGM was ∼US$16 million or about eight US cents per person affected (Herren & Neuenschwander, 1991). This well-studied system provides important insights applicable to *ex ante* analyses of future biological control efforts, especially given the current capacity to project PBDM results over wide geographic areas. In our cassava example, *ex ante* analyses would have given the correct predictions concerning efficacy of the natural enemies of both CM and CGM across Africa..

Specifically, our *post hoc* physiologically based demographic model (PBDM) analysis of the cassava system captured the effects on yield and the observed distribution and relative abundance of CM and CGM and their natural enemies across Africa astonishingly well; independent of species distribution records. It showed that natural enemies, rather than weather or resistant varieties caused the decline of CM and CGM populations. The analysis explained why the parasitoid *A. lopezi* introduced from a small region in the Rio de la Plata Valley in South America quickly established with immediate high impact on CM across the ecological zones of Africa (Fig. 4A; Neuenschwander, 2001, 2004; Hammond & Neuenschwander, 1990). The analysis also explains that if only *A. diversicornis* had been introduced, only partial control of CM would have occurred, resulting in a biological control failure (see Fig. 4B). The model explained the role of the biology of the parasitoids in the control of CM and their interactions (see Gutierrez, Neuenschwander & Alphen 1993).

The search for predators of CGM was focused using a climate matching program developed at CGIAR/CIAT in Colombia (Yaninek & Bellotti, 1987) enabling geographic concentration of the search for adapted predators and pathogens in Brazil. Eleven species of mite predator were introduce to Africa, but only *A. manihoti* and *T. aripo* established with *T. aripo* becoming the dominant predator in the field (Yaninek *et al*., 1993). The analysis explains that if only *A. manihoti* had been introduced, only partial control of CGM would have occurred resulting in another failure. The model also explains as observed that populations of CM and CGM may still develop in ∼ten percent of fields, especially in dry marginal areas (Fig. 6B and 8F respectively).

An excellent *ex ante* PBDM analysis of natural enemy efficacy is the coffee (*Coffea arabica* L.) system that explained why several introduced parasitoids of coffee berry borer (CBB, *Hypothenemus hampei* (Ferrari)) in the Americas would fail (see Cure *et al*., 2020). Further, Abram *et al*. (2020) used a simple model to posit that egg parasitoids of the invasive Asian brown marmorated stink bug (BMSB, *Halyomorpha halys* Stål) were unlikely to control the pest across its vast Palearctic range, while a tri-trophic PBDM system explained why, and further posited *ex ante* that tachinid parasitoids of the large nymphal and adult stages would have an important role in suppressing BMSB (Gutierrez *et al*., 2023). The failed control of yellow starthistle (*Centaurea solstitialis* L) in North America by introduced seed head insects was explained and *ex ante* analysis posits that control would be greatly enhanced by the introduction of the rosette weevil (*Ceratapion basicorne* (Illiger)) that kills whole plants and/or greatly reduces seed production in survivors (Gutierrez et al. 2017). These examples (see also Gutierrez et al. 2025a) suggest that pre-release *ex ante* analyses of biological control agents using well parameterized weather driven PBDMs would be advantageous, cost effective and would complement species distribution (e.g., Elith & Leathwick, 2009) and genomic analyses (Williams *et al*., 1994).

But PBDM development is perceived difficult (Barlow, 1999), the process is straightforward Gutierrez & Ponti, 2013; Gutierrez *et al*. (2025a). The bottle neck, however, will often be developing the appropriate biological data to parameterize PBDMs. Such studies are often deemed mere technical work and is not overly appreciated or rewarded professionally, and yet such studies are key to answering larger ecological and economic/social questions (Gutierrez *et al*., 2015; 2025b). On a positive note, collaborative research is in progress to develop software to enable non-experts to develop PBDMs (Simmons *et al*. 2025).

A pest problem currently constraining cassava production in Africa that begs attention is whitefly (*Bemisia tabaci* (Glenn.)) that vectors mosaic disease (ACMD) and cassava brown streak disease (CBSD). Control of *B. tabaci* in resource poor cassava production in Africa requires control methods that are low-cost, effective, sustainable, and readily disseminated, such as traditional host-plant resistance and biological control (Legg *et al*., 2015). This problem requires holistic agro-eco-social analyses (*cf*. Nagel, 2012), and *ex ante* PBDM analyses of the complicated biology of whitefly parasitoids (Mills and Gutierrez 1996) must play an important role.

Last, while the PBDM approach tends toward reductionism, from a theoretical standpoint it has roots in thermodynamics with an explicative basis for population interactions provided by bioeconomic supply-demand principles common to all economies including human economies (see Gutierrez 1992; Regev *et al*., 1998; Gutierrez and Regev 2006). These properties allow PBDMs to play an important role in holistic agroecosystem analyses including *ex ante* evaluation of natural control agent efficacy, and demonstrating this was raison d’être of this study.

## Declarations

### Ethics approval and consent to participate

not applicable.

### Consent for publication

Not applicable.

### Availability of data and material

All the data used is available in the cited references.

### Authors’ contributions

All authors contributed equally.

## Acknowledgements

We are grateful to the international network maintaining the Geographic Resources Analysis Support System (GRASS) for making the GIS software available to the scientific community.

