## Supplemental Materials for "*Ex ante* analyses can predict natural enemy efficacy in biological control"

> *An important transformation in ecological research occurred when system thinking shifted the attention of ecologists from objects to relationships and from structures to processes - from objects to networks of relationships embedded in larger networks - and from fundamental structures affected by forces and mechanisms to the recognition that every structure is a manifestation of underlying processes* (Capra & Luisi, 2014).

Our analyses delve into the details of the biology of the cassava system, but the goal is not a one-to-one description of the biology of the system, rather the goal is to capture sufficiently the weather-driven relationships underpinning the system sufficiently to make prospective model predictions independent of time and place. Interested readers should review the cited papers for full details of the cassava system model.

**Cassava mealybug and its natural enemies**

The model is mostly based on laboratory studies (Löhr et al., 1988; Neuenschwander & Madojemu, 1986; Neuenschwander et al., 1986), and describes the distribution and impact of the cassava mealybug (*Phenacoccus manihoti* Matile-Ferrero (Hemiptera, Pseudococcidae))(CM) and *Anagyrus. lopezi* DeSantis (Hymenoptera, Encyrtidae) astonishingly well as reported in surveys across Africa (Boussienguet et al., 1991; Chakupurakal et al., 1994; Neuenschwander et al., 1986; Neuenschwander, Hammond, et al., 1987; Neuenschwander, Hennessey, et al., 1987). About 10% of all fields develop damaging CM populations depending on the season and how long *A. lopezi* has been established, with outbreaks mostly confined to sandy, un-mulched soils. Mulching improved the condition of the plant and increased CM size and hence the performance of *A. lopezi* developing on them (Schulthess et al., 1997). Indigenous (and introduced) predators as well as non-specific hyperparasitoids become locally abundant only under conditions of high mealybug infestation, but do not contribute measurably to CM control (Goergen & Neuenschwander, 1992; Neuenschwander, Hammond, et al., 1987; Neuenschwander & Hammond, 1988; Neuenschwander & Sullivan, 1987; Stäubli Dreyer, Baumgärtner, et al., 1997; Stäubli Dreyer, Neuenschwander, et al., 1997). Predatory coccinellids removed about 10% of high peak CM populations (Gutierrez et al., 1993) and hyperparasitoids behaved strictly in a density dependent manner, and hence they are mostly absent at the common low CM densities (Neuenschwander & Hammond, 1988). For this reason, they were not included in this study.

Laboratory studies indicate that *A. lopezi* out-competes *A. diversicornis* (Howard) in cases of multiple parasitism because it has a higher search efficiency, attacks smaller CM, and has a higher female-biased sex ratio under adverse conditions that produce smaller mealybugs (Pijls et al., 1995). These attributes enabled *A. lopezi* to dominate across Africa as shown by the model. The dominant role of *A. lopezi* was further documented in exclusion experiments (Cudjoe et al., 1992, 1993; Neuenschwander et al., 1986) and reconfirmed when it was released in Southeast Asia for control of CM (Wyckhuys et al., 2018).

**Cassava green mite and its natural enemies**

As with CM, the model parameters for the cassava green mite (*Mononychellus tanajoa* (Bondar), (Trombidiformes, Tetranychidae)) (CGM) and its natural enemies are primarily from laboratory studies (see Gutierrez, Yaninek, et al., 1988; Gutierrez et al., 1999 for relevant citations) and the simulations reflect the known distributions and impact well. In its native range in the Neotropics, CGM populations are maintained under natural control by a combination of local predators, pathogens, and weather (e.g., rainfall mortality), especially where cassava has been locally selected and cultivated in a traditional manner (Yaseen & Bennett, 1977; Bellotti & Schoonhoven, 1978; Samways, 1979; Elliot et al., 2000).

After the accidental introduction of CGM in Africa, local natural enemies, largely opportunistic generalist predators like coccinellids in the genus *Stethorus* and staphylinids in the genus *Holobus* (= *Oligota*) ( Yaninek & Herren, 1988) and endemic fungal pathogens never associated with epizootics (Yaninek et al., 1996) were present, but were not effective biological control agents. Yield losses following the invasion ranged from 13 to 80% (Lyon, W. F., 1974; Nyiira, 1976; Shukla, 1976; Ndayiragije, 1984; Yaninek et al., 1990). CGM feed on the under surface of young leaves killing leaf cells reducing the photosynthetically active leaf area with densities varying with plant drought stress, leaf age and cultivar (Yaninek, Gutierrez, et al., 1989; Yaninek, Herren, et al., 1989; Yaninek et al., 1990). Yaninek , Gutierrez *et al*. (1989) found that at 24°C, average longievity of CGM adult female in Africa was ~ 32 days and produced ~58 eggs over ~16 days, while Moraes *et al.* (1995) under similar conditions in the Neotropics found that at 24°C CGM adult female lived an average of ~ 30 days and produced ~42 eggs over ~15 days.

The interaction biology of CGM and its native predators is complicated. All the released mite predators (Mesostigmata, Phytoseiidae) that established in Africa were selected for introduction based on agrometeorological matching criteria (Yaninek & Bellotti, 1987). Subsequent studies revealed that *Typhlodromalus aripo* De Leon, the apparently least effective predator because it was less voracious and slower in population increase than *Amblydromalus manihoti* Moraes and *Neoseiulus. idaeus* Denmark & Muma, proved to be the best biological control agent for *M. tanajoa* in Africa.

***Typhlodromalus aripo***: *T. aripo* establishment, dispersal and persistence on cassava (Onzo et al., 2003) is enhance by it efficiently located prey, and persists at low prey densities by using pollen, and cassava extrafloral exudates as an alternative food sources (Gnanvossou et al., 2001). It normally inhabits a refuge located in-between the leaf primordia at the apex of the cassava plant that is thought to provide protection ( a refuge) against abiotic factors and intraguild predation (Onzo et al., 2010). *T. aripo* shows a pronounced diurnal within-plant change in distribution and elicit an avoidance response in mobile CGM stages causing them to disperse and becoming less abundant on the first twenty leaves below the apex during the night compared to the same leaves in late afternoon (Onzo et al., 2003). *T. aripo* also displays a marked attraction to odors emitted from either CGM-infested apices or infested young leaves over infested old leaves but showed no preference for odors from apices versus young leaves (Gnanvossou, Hanna, et al., 2003). Under high prey densities, the average longevity of *T. aripo* adults is about twice that of *A. manihoti*, but as prey densities fall, reproduction in *T. aripo* declines but its longevity is double that of fully fed adults (see Magalhães et al., 2003). This occurs because *T. aripo* feeds on pollen enabling it to extend its longevity when prey is scarce, but with reduced fecundity (Gnanvossou et al., 2005). *T. aripo* exhibits a trade-off in the metabolic rate: it has a low metabolic rate which allows long survival periods but slows oviposition and development (*cf.* Magalhães et al., 2003). This trade-off in biology is captured in the *T. aripo* model by scaling the daily change in physiological time ($^{s}\Delta x(T(t))$, see below) and fecundity by the food supply/demand ratio (S/D): i.e., that decreases both the developmental rate and reduces fecundity. However, under abundant prey, the functional response of *T. aripo* females to CGM egg density is type II reaching an asymptote at ~175 CGM eggs attacked/day under experimental conditions (Cuellar et al., 2001). However, its numerical response is quite low reaching a maximum of ~1.2 to 1.4 eggs/day at 100 CGM eggs attacked/day (Cuellar et al., 2001; Gnanvossou, Yaninek, et al., 2003). Interestingly, one *T aripo* egg is produced at 22 prey eggs attacked, and thereafter, the reproductive rate levels off at ~1.3 eggs/day despite a high attack rate. This suggests that at high prey densities, *T. aripo* can kill four-fold more prey eggs (or egg equivalents) than required for reproduction (see Cuellar et al., 2001). Further, Mutisya (2014) found the thermal limits for *T. aripo,* are 11.4°C for development with oviposition occurring between 12°C and 33°C with the optimum at about 27°C. High fecundity occurs in the range 25 to 100% RH with an optimum at ~75% RH with an adult longevity of ~426 dd.

***Amblydromalus manihoti***: In sharp contrast, *A. manihoti* is found only on fully developed cassava leaves and preferentially forages in the middle of the foliage (Bonato et al., 1999). It is a generalist mite predator and does not discriminate between volatiles from different infested cassava leaves (Gnanvossou, Hanna, et al., 2003) lowering its search rate on cassava. It has a high metabolic rate; rapidly consumes prey that it converts to eggs at the expense of lower survival and longevity (Gnanvossou, Yaninek, et al., 2003). Furthermore, *A. manihoti* does not feed on pollen and readily migrates when prey populations on cassava are low (Yaninek et al., 1998). The lower thermal threshold of *A. manihoti* is 6.64°C for development, and oviposition occurs between 6.64°C and 31°C with an optimum at about 19°C above 40% RH. The thermal parameters for *A. manihoti* are those often associated with a temperate species. Artificially increasing the lower thermal threshold to 11°C in the model did not measurably change the conclusions as the other aspects of the biology matter.

### System model overview

#### Cassava model

The original cassava system model was a canopy model composed of age-mass structured population sub-models of leaf, stem, root (tuber), pests, and natural enemies/pathogens (Gutierrez, Wermelinger, et al., 1988; Gutierrez, Neuenschwander, et al., 1988; Gutierrez, Yaninek, et al., 1988; Gutierrez et al., 1993). The canopy model was extended to be a meta-population model that could be up to 100 plants evenly or randomly spaced with each plant having different areas for growth and each potentially having the full complement of arthropod species in our study (Gutierrez et al., 1999). However, for daily computations over a ten-year period on a laptop computer, a system of ten randomly spaced plants was used in our study.

Plant models are usually constructed as metabolic pool models (MP) that capture the effects of per capita biomass/energy demand (D) and realized acquisition via search (S) and allocation to respiration, growth, and reproduction. Note that S<D. Specifically, plants search for light, water, nutrients to produce photosynthate that is allocated to plant subunit growth and to reproduction (*cf.* de Wit & Goudriaan, 1978). Gutierrez and Baumgärtner (1984) used MP notions to develop PBDM models for growth, development, and reproduction of arthropods. Most of the species in the cassava system model are MP based (Table 1).

Table 1. Type of model used for each stage/species.

| species | all stages | eggs | immatures | adults |
|---|---|---|---|---|
| cassava | MP | | | |
| CM | MP | | | |
| *A. lopezi* | MP | MP | BDF | |
| *A. diversicornis* | | MP | MP | BDF |
| CGM | MP | | | |
| *T. aripo* | MP | | | |
| *A. manihoti* | MP | | | |
| pathogens | BDF* | | | |

*A function of daily rainfall.

Biodemographic functions (BDF) can also be developed to characterize developmental rates and birth and death rates of species (*cf.* Gilbert et al., 1976). These vital rates are usually estimated from age-specific life table studies conducted under an array of temperatures and other conditions (Gutierrez et al., 2021; Gutierrez & Ponti, 2013) and are the outcomes over the life cycle of a cohort of organisms of how they acquired and allocated energy, survived, and reproduced under the experimental conditions – i.e., metabolic pool processes. These functions can also be estimated from field ecological studies (Maurer & Baumgärtner, 1992, 1994; Fouque et al., 1992; Fouque & Baumgärtner, 1996; Dreyer & Baumgärtner, 1995).

The BDFs commonly used to estimate the effects of abiotic variables on species developmental, birth and death rates in age-structured, weather-driven BDF models are depicted in a stylized manner in Fig. 2 in the text. Other BDFs can be developed to accommodate additional aspects of the biology of a species.

**Developmental rate:** Time and age for all species are in physiological time units that differ among the species. An improved component of the system model is a nonlinear developmental rate model for cassava where *r(T)* was estimated from data in Keating and Evenson (1979).

$$r(T) = 1/d(T) = \frac{a(T-\theta_L)}{(1+b^{T-\theta_I})} = \frac{0.0095(T-14.85)}{(1+1.6^{T-33.55})} \quad (1)$$

*d(T)* is development time in days at temperature *T*, with fitted constants a, b, $r(T)=0$ at $T \leq \theta_L$=14.85°C, the maximum developmental rate occurs at $\theta_I$ ~33.55°C, and $\theta_U$= 41°C is the upper thermal threshold. Due to data limitations, linear developmental rate models (i.e., ${}^{s}r(T) = a + bT$ ) were used for the other species (left superscript s), with ${}^{s}r(T)=0$ at $T \leq \theta_L$. The physiological developmental time constant (${}^{s}\Delta$) in degree days (*dd*) for each life stage of a species was estimated in the linear range of favorable temperatures as ${}^{s}\Delta = d(T) \times (T - {}^{s}\theta_L)$ (Campbell et al., 1974). The daily increment of physiological time at time *t* and temperature *T* is ${}^{s}\Delta x(T(t)) = {}^{s}r(T(t))\,{}^{s}\Delta$ . Hence, on average a cohort of individuals initiated at time $t_0$ completes development when $\int_{t_0}^{t} {}^{s}r(T(t))dt = 1$ in continuous form, or $\sum_{t_0}^{t} {}^{s}\Delta x(T(t)) = {}^{s}\Delta$ in *dd* units in our discrete time models. Further, each species (and their stages) may age on different temperature-dependent time scales. As noted above, at low prey density, *T. aripo* increasingly feeds on pollen that increases longevity but decreases fecundity. This biology is incorporated in the model by using the success rate of *T. aripo* feeding on CGM prey (S/D<1) is used to scale ${}^{Ta}\Delta x(T(t))^{Ta}(S/D)$ (i.e., slowing the developmental rate) and population level fecundity (*E*) (see below).

**Resource acquisition and allocation**

All organisms are consumers. Stage-specific resource acquisition (S) is demand-driven (D) and may be in mass or number units. Population resource demands (D) for a species/stage (left superscript *s*) (see Gutierrez 1992, 1996) are computed as

$${}^{s}D = \sum_{i=2}^{7} {}^{s}N_i \, {}^{s}D_i \, {}^{s}\Delta x(T(t))$$ **,** where ${}^{s}D_i$ is the per capita demand $dd^{-1}$ at age *i*, ${}^{s}N_i$ is the number of individuals of age *i*, and $\Delta x(T(t)$ is the *dd* at time *t* and temperature *T*. Population-level resource supply (S) obtained by species/stage (s) (text Fig. 2B) is computed in the predator form of the functional response model as

$${}^{s}S = {}^{s}D(1 - e^{\frac{-\,{}^{s}\alpha\, prey}{{}^{s}D}}) \qquad \{2\}$$

where $a$ is the consumer search rate.

Using the MP approach, the demand for a plant ( ${}^{plant}D$ ) or say a mealybug ${}^{CM}D$ ) in mass units would be the total of all age and temperature dependent subunit growth and reproduction demand ( $D_G + D_R$ ) plus respiration costs ( $vM$ ) corrected for assimilation efficiency ( $(1-\beta)$ ).

$$\begin{aligned} {}^{plant}D &= (D_G + D_R + vM) / (1-\beta) \\ {}^{plant}S &= {}^{plant}D(1 - e^{\frac{-\alpha \text{ resource}}{{}^{plant}D}}) \end{aligned} \qquad \{3\}$$

This form assumes the resource attacked is not available to other consumers (e.g., light captured by leaves, prey consumed by a predator).

However, some resources may be attacked more than once. For example, a parasitoid female attacks whole individuals that may also be attacked more than once by the same or other parasitoids (super and multiple parasitism. In this case, ${}^{s}D$ is the per capita number of *hosts* an adult female parasitoid can attack (i.e., its egg load) and ${}^{s}D = \sum_{i=2}^{7} {}^{s}N_i \, {}^{s}D_i \, {}^{s}\Delta x(T(t))$ is the population attack demand. Using the parasitoid form of the model,

$${}^{s}S = hosts \times proportion\ attacked = hosts(1 - e^{\frac{-\,{}^{s}D}{H}(1-e^{\frac{-\,{}^{s}\alpha\, H}{{}^{s}D}})}) \qquad \{4\}$$

where $a$ is the parasitoid search rate and *H* is the available hosts. In all cases $0 < {}^{s}(S/D) < 1$.

In a more realistic model (eqn. 5), realized demand by a parasitoid population (${}^{s}D$) is the per capita fecundity ( ${}^{s}f(x)^{-dd}$ ) at age *x* at optimal temperature ($T_{opt}$) (see text Fig. 2C), ${}^{s}sr$ is the sex ratio and ${}^{s}\phi_T$ scales the demand for temperature effects in a time temperature varying environment.

$${}^{s}D = {}^{s}\phi_T \cdot {}^{s}sr \cdot \int {}^{s,adults}N \cdot {}^{s}f(x)^{-dd}dx \;\; {}^{s}\Delta x(T(t)) \qquad \{5\}$$

Variations of the above biology occur for plants and predacious organisms, but by analogy all of them can be accommodated by these simple functions (see Gutierrez, 1996).

**Cassava mealybug life stages and parameters**

The mealybug model is a mass-age structured MP model. Age windows for the CM stages in *dd*>14.6°C are:

```
0          114            182          306          371                 890dd
|--eggs---|--crawlers--|--larv2, 3--|-- adults --|--adults ovip---|.
```

The total eggs produced per day by the population using a BDF is:

$${}^{CM}E = {}^{CM}sr\,{}^{CM}\phi(T)\,{}^{CM}(S/D)\sum_{i=1}^{k_{CM}}((7.1/(1+1.325^{i}))CM_i$$

where ${}^{CM}\phi(T) = (T,\ 18^{\circ}C,\ 35^{\circ}C)$, ${}^{CM}(S/D)$ is the supply/demand ratio, and the effects of temperature enter as a symmetrical scalar function of temperature

$$0 \le {}^{CM}f(T) = 1.0 - (T - \theta_L - A)/A)^2 \le 1 \text{ where A} = (\theta_L - \theta_U)/2.0 \text{ (see text Fig. 2D).}$$

The notation ${}^{CM}\phi(T)$ is used for CM, but a similar notation with left superscript is used to denote the same functional form for the other species (e.g., ${}^{Al}\phi(T)$, ${}^{Ad}\phi(T)$, ${}^{CGM}\phi(T)$, etc.). Other similar scalar functions can be developed and their product used to scale fecundity (e.g., ${}^{CM}\phi(Nitrogen) \times {}^{CM}\phi(RH) \times {}^{CM}\phi(pH)$) or other biological processes for this and other systems.

Rainfall/fungal mortality on CM is a simple empirical model:

$$0 \le {}^{CM}\mu_{P^+} = 0.45(1 - e^{-0.025\,precip}) \le 1.$$

**Cassava mealybug natural enemies**

Prior studies (see above) indicated *A. lopezi* out competes *A. diversicornis* in cases of multiple parasitism, it has a higher search efficiency, attacks smaller CM, and has a higher female-biased sex ratio under adverse conditions that produce smaller mealybugs (see Gutierrez et al., 1993). These attributes enabled *A. lopezi* to dominate across Africa.

***A. lopezi* life stages and parameters**

age := transit times for *A. lopezi* in *dd* >13.5°C

```
age1-------------age2--------age3------------------------------age4
0                100         186                               366dd
 |--eggs ---larva-|--pupa----- | --------------------adults ---------|
```

$$^{Al}D = {}^{Al}sr\,{}^{Al}\phi(T)\{{}^{Al}ovip(1+{}^{Al}hf)\int {}^{Al}Adlt\;dx\}\,{}^{Al}\Delta x$$

where $^{Al}hf$ =0.2 corrects for host feeding, the symetrical concave function $0 \le {}^{Al}\phi(T) = (T,\ 13.5,\ 34) \le 1$ corrects for temperature limits of 13.5°C and 34°C, $^{Al}ovip = 0.9$ is fecundity per *dd*, $^{Al}sr \geq 0.5$♀♀ is the sex ratio, and $^{Al}\Delta x$ is the change in age/time. Potential hosts ($^{Al}H$) for *A. lopezi* weighted for *CM* stage preferences and multiple and super parasitism (preference for parasitoid egg-larval stages of Al[1] and Ad[1]) are computed as follows:

$$^{Al}H = 0.328\ CM[3] + 0.514\ CM[4] + 1.000\ CM[5] + 1.000\ CM[6] + 0.25\ Al[1] + 0.25\ Ad[1]$$

where $^{Al}H$ replaces $H$ in the parasitoid functional response eqn. 4.

$0 < {}^{Al}\mu = \left(1 - e^{\frac{-{}^{Al}D}{{}^{Al}H}\left(1-e^{\frac{-{}^{Al}\alpha\ {}^{AL}H}{{}^{Al}D}}\right)}\right) < 1$ and $^{Al}a = 0.65$ is the parasitoid search rate with $^{Al}Na = {}^{Al}H\,{}^{Al}\mu$ being the number of hosts attacked. The computations for attack rate ($0 < {}^{s}\mu \le 1$) of CM and parasitoid host stages attacked by *A. lopezi* corrected for preference as follows:

$$^{CM}\mu[3] = {}^{Al}Na \times (0.328\ CM[3]/{}^{Al}H)/CM[3]$$

$$^{CM}\mu[4] = {}^{Al}Na \times (0.514\ CM[4]/{}^{Al}H)/CM[4]$$

$$^{CM}\mu[5] = {}^{Al}Na \times (CM[5]\ /{}^{Al}H)/CM[5]$$

$$^{CM}\mu[6] = {}^{Al}Na \times (CM[6]\ /{}^{Al}H)/CM[6]$$

$$^{Al}\mu[1] = {}^{Al}Na \times (0.250\ Al[1]/{}^{Al}H)/Al[1]$$

$$^{Ad}\mu[1] = {}^{Al}Na \times (0.250\ Ad[1]/{}^{Al}H)/Ad[1]$$

Similar computations occur for *A. diversicornis,* and if both parasitoids are present, the attacked hosts are distributed as illustrated by the Venn diagram below, noting that in cases of multiple parasitism, *A. lopezi* wins (i.e., the shaded area).

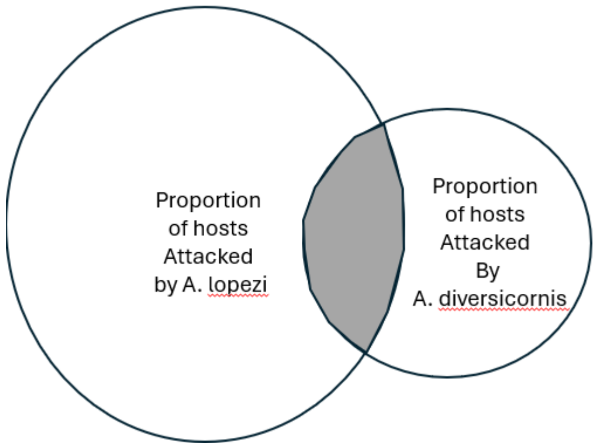

#### *A. diversicornus* life stages and parameters

age := transit times for *A. diversicornus* in *dd* >13.5°C

```
(*  age1----------------age2-------age3------------------------------age4
    0                   105           195                            380dd
    |--eggs -------larva-|--pupa-----| --------------------adults-------|  *)
```

Potential hosts ($^{Al}H$) for *A. diversicornis* weighted for *CM* stage preferences (i.e., stages [3-6]) plus preference of parasitoid egg-larval stage [1] resulting in multiple and super parasitism:

$$^{Ad}H = 0.160\ CM[3] + 0.540\ CM[4] + 1.000\ CM[5] + 0.50\ CM[6] + 0.25\ Al + 0.25\ Ad,$$

and the demand for hosts is

$$^{Ad}D = {}^{Ad}sr\, {}^{Al}\phi(T)[{}^{Ad}ovip(1 + {}^{Ad}hf)\int {}^{Ad}adlt\ dx]^{Ad}\Delta x$$ with parameters

$^{Ad}\phi(T) = (T,\ 13.5°C,\ 34°C)$, $^{Ad}ovip$ = 0.9/dd, and $^{Ad}sr$ = 0.5♀. $^{Al}\alpha$ = 0.6 is the search rate in eqn. 4.

#### Cassava green mite life stages and parameters

age := transit times for *CGM* in *dd* >14.65°C

```
(* cassava green mite
0                          86        98                              418dd
 |--------immatures--------|-preova-|--------adults ovip---------|
*)
```

Reproduction in CGM is computed as

$$^{CGM}E = {}^{GM}sr\, {}^{CGM}\phi(T)\, {}^{CGM}(S/D)\sum_{1=1}^{k} 1.8i/(1+1.15^{a})CM(i),$$

where $^{GM}$sr = 0.8 female biased,

$0 \le {}^{CGM}\phi(T) = \left(-0.0164T^{4} + 1.7381T^{3} - 69.118T^{2} + 1224.7T - 8104\right)/62.8 \le 1$ is the normalized left skewed effect of temperature (Yaninek et al., 1989),

$0 \le {}^{GM}\mu_{P^{+}} = 1.0 - e^{-0.0185\,precip} < 1$ is the net rainfall/fungal mortality rate,

and 0.0 ≤ $^{CGM}\mu(T)$ = $(0.000012T^{4} - 0.001187T^{3} + 0.042353T^{2} - 0.666530T + 3.938533) \le 1$ is the temperature-dependent mortality rate (e.g., text Fig. 2E) where *T*=*Tmean* (see text Fig. 2E).

#### *T. aripo* life stages and parameters

age := transit times for *T. aripo* in *dd* > $^{Ta}\theta_L$ =11.4°C ( Gnanvossou et al., 2003)

```
(* T. aripo
 0    28.6   42.2    58.48    74.8   110.2           350.0          396.0dd
 |-ova-|-larv-|-proto-|-deuto-|-preov-|--ovip-------|--post ovip --|
 0      1       2          3          4         5                  6                 7
*)
```

Predator age classes [1-7] μg demand for prey in prey egg equivalents/dd at 25°C.

$^{Ta}$Dem[1]:=0.0

$^{Ta}$ Dem[2]:=0.24μg {micrograms per dd equals 9.4 prey eggs over the laval stage }

$^{Ta}$ Dem[3]:=0.30μg { 15.6 eggs - protonymph stage}

$^{Ta}$ Dem[4]:=0.56μg { 25 eggs - deutonymph stage}

$^{Ta}$ Dem[5]:=0.54μg { 89 eggs - preoviposition female adult}

$^{Ta}$ Dem[6]:=0.58μg {191 eggs over female oviposition period}

$^{Ta}$ Dem[7]:=0.45μg { 95 eggs post-oviposition period}

$^{Ta}$ fecundity:= 0.084 eggs/ *dd> 11.4℃* (Yaninek estimate 11/21/24 see MUTISYA et al. 2014)

$^{Ta}$sr= 0.66 {2 females:male}

$^{Ta}$RHlim< 25.0% (limiting RH)

$^{Ta}$SD< 0.5 {pollen feeding begins and enables longer survival and reduced reproduction}

($^{Ta}\alpha$ ):= 1.2 {search rate}

$0 \leq {}^{Ta}\phi(T) = (T,\ 11.4°C,\ 35.4°C) \leq 1$ (temperature scalar with limits 11.4-35.4°C; Mutisya et al., 2014)

$^{Ta}\mu(RH)$ = min(0.99, 1-$exp^{-2.0/(RHmean-RHlim)}$) is desiccation mortality rate where RHlim= 10%.

***A. manihoti* life stages and parameters**

age := transit times for *A. manihoti* in *dd* > $^{Aml}\theta_L$ =6.64°C

```
(*
A. manihoti substage age intervals in degree days >  6.64°C)
    0    39.06  61.4    81.8     98.6     120.9      254.8          288.30
    |-ova-|-larv-|-proto-|-deuto-|-preov-|--ovip-------|--post ovip --|
    0      1       2         3         4         5                6                7
*)
```

Predator age classes [1-7] μg demand for prey in prey egg equivalents/dd at 25°C{it attacks its own)}

$^{Am}$Dem[1] = 0.0

$^{Am}$Dem[2] = 0.22μg { 1 μg per dd}{ 16.0 eggs - laval stage }

$^{Am}$Dem[3] = 0.21μg { 18 eggs - protonymph stage }

$^{Am}$mDem[4] = 0.26μg { 23 eggs - deutonymph stage }

$^{Am}$Dem[5] = 0.41μg { 98 eggs - preoviposition }

$^{Am}$Dem[6] = 0.37μg {257 - oviposition period }

$^{Am}$Dem[7] = 0.27μg { 63 - post-oviposition at 25C}

$^{Am}$fecundity = 0.18 eggs/dd>6.64°C

$^{Am}$sr = 0.66 {2 females:male}

$^{Am}$RHlim = 40%

$0 \leq {}^{Am}SD_{min} \leq 1$ {no pollen effect for *A. manihoti*}

$^{Am}\alpha =$ 0.8 {search rate < random}

$^{Am}\phi(T) = (T,\ 6.64°C,\ 32.8°C)$ (temperature scalar with limits 6.64 - 32.8°C

Desiccation mortality where VPD is vapor pressure deficit (see Mégevand, 1997)

$^{Am}\mu(T,Rh)$ = max (0, min (0.99, 1.0 - 0.1658 VPD + 1.1054) where

$$VPD = 610.7 \cdot (10^{7.5T/(237.3+T)})(1 - Rh/100)/1000$$

and $T$ is mean temperature °C.

**Distributed maturation time population dynamics model**

The models used herein are time varying life tables (TVLT), the parameter of which may be MP or BDF based (see above). Suitable population dynamics models used to capture the dynamics of interacting species were reviewed by Gutierrez (1996), Di Cola *et al.* (1999), and Buffoni and Pasquali (2007). For simplicity, we use the discrete form of the time-invariant distributed-maturation time demographic models (Abkin & Wolf, 1976; Manetsch, 1976) parameterized using the MP and/or BDF biology. The time-varying form of the model (Vansickle, 1977) is appropriate where the developmental time of the species and substages change over time in response to various factors (e.g., nutrition).

In the model, the dynamics of a life stage $s$ of average developmental time ${}^s\Delta$ having $i$=1, 2,...,${}^sk$ age classes can be viewed as composed of ${}^sk$ dynamics equations (eqn. 5; Gutierrez, 1996; Severini et al., 2005). Using the notation of Di Cola *et al.* (1999, page 523), the $i^{th}$ age class of stage $s$ is modeled as follows:

$$\frac{d\,{}^sN_i}{dt} = \frac{{}^sk \cdot {}^s\Delta x}{{}^s\Delta}\left[{}^sN_{i-1}(t) - {}^sN_i(t)\right] - {}^s\mu_i(t)\,{}^sN_i(t). \qquad \text{[A1i]}$$

In terms of flux, ${}^sn_i(t) = {}^sN_i(t)\,{}^s\nu_i(t)$ where ${}^s\nu_i(t) = \frac{{}^sk}{{}^s\Delta}\Delta x(t)$, and

$$\frac{d}{dt}\left[\frac{{}^s\Delta\,{}^sn_i(t)}{{}^sk}\right] = {}^sn_{i-1}(t) - {}^sn_i(t) - {}^s\mu_i(t)\,{}^sn_i(t)\frac{{}^s\Delta}{{}^sk}. \qquad \text{[A1ii]}$$

Absent mortality, the theoretical distribution of cohort developmental times of stage $s$ may be estimated by Erlang parameter ${}^sk = {}^s\Delta^2 / {}^s\sigma^2$, where $\sigma^2$ is the variance of ${}^s\Delta$ (i.e., theoretically, a cohort entering a stage will have Erlang distributed maturation times centered on ${}^s\Delta$). Appropriate data were unavailable, an hence a value of $k$ = 50 was used for all of the species.

The forcing variable is temperature ($T$), with chronological time ($t$) in days ($d$) that from the perspective of poikilotherm species is of variable length in physiological time units (i.e., $0 \leq {}^s\Delta x(T(t))$ in degree days ($dd$)), or proportional development (${}^sr(T(t))$). The state variable ${}^sN_i(t)$ is the density of the $i^{th}$ age class (mass or numbers), and ${}^{s}\mu_i(t)$ is the proportional age-specific net loss rate (losses and gains) due to temperature, net immigration, growth in mass dynamics models, and other factors during ${}^{s}\Delta x(T(t))$ (Gutierrez, 1996). For computational efficiency in the model, the components of ${}^{s}\mu_i(t)$ are computed and applied after aging. The total density of a life stage *s* is ${}^{s}N(t) = \sum_{i=1}^{k} {}^{s}N_i(t)$. New individuals enter the first age class of a stage 1, flows occur via aging between age classes and between stages, and surviving adults exit as deaths from maximum age ($i = {}^{A}k$).

Variations of ${}^{s}\mu_i(t)$ in this model easily facilitate capturing the biology of any species. The numerical solution for eqn. A1ii is found in Abkin and Wolf (1976), and as implemented here in Gutierrez (1996, pages 157-159).
